# Both T and B cells contribute to dysregulated activation and differentiation of CD4^+^ T cells in Activated PI3K delta syndrome 1

**DOI:** 10.1101/2024.08.04.606503

**Authors:** Julia Bier, Anthony Lau, Katherine JL Jackson, Stephanie Ruiz-Diaz, Robert Brink, Stuart G Tangye, Elissa K Deenick

## Abstract

Activated PI3K delta syndrome 1 (APDS1) is caused by a heterozygous germline gain-of-function (GOF) variants in *PIK3CD* which encodes the p110δ catalytic subunit of phosphoinositide 3-kinase (PI3K). APDS1 patients display a broad range of clinical manifestations and perturbations in cellular phenotype. One of the most striking features is the dysregulation of the T cell compartment characterised by an increase in memory T cells, including Tfh cells, and a concomitant decrease in naïve T cells. We have previously shown that many of these changes in T cell populations were T cell extrinsic and were also induced in WT T cells that developed in the presence of PI3K GOF cells. Here we dissected the drivers of dysregulated T cell activation using a mouse model of APDS1. This revealed PI3K GOF macrophages and DCs made little contribution to the aberrant T cell activation. Instead, the loss of naïve T cells was mostly driven extrinsically by PI3K GOF T cells, while the increase in Tfh cells was mediated by dysregulated PI3K GOF B cells. Surprisingly, despite previous reports of increased PI3K driving dysregulated inflammatory Tregs, we saw no evidence for *Pik3cd* GOF Tregs acquiring an inflammatory phenotype and driving T cell activation. These studies provide new insights into the clinical phenotype of patients with APDS1 and new understanding of the role of PI3K in immune cells.

## Introduction

Activated PI3K delta syndrome 1 (APDS1) is an immune dysregulatory condition caused by heterozygous germline gain-of-function (GOF) variants in *PIK3CD* [1–3]. *PIK3CD* encodes p110δ, a catalytic subunit of phosphoinositide 3-kinase (PI3K), which is primarily expressed in lymphoid cells [4]. PI3K is activated downstream of many key immune receptors, where it promotes phosphorylation of phosphatidylinositol-(4,5)-biphosphate (PIP_2_) to phosphatidylinositol-(3,4,5)-trisphosphate (PIP_3_), leading to the activation of multiple downstream molecules, including AKT and mTOR [1, 4]. The PI3K signalling pathway regulates multiple functions in immune cells, including cell development, differentiation, metabolism and activation [4]. Thus, it is important that PI3K activity is tightly controlled. Indeed, multiple layers of regulation exist including the phosphatases PTEN and SHIP, which dephosphorylate PIP_3_ and thus counteract the action of PI3K [4, 5].

The *PIK3CD* GOF variants in APDS1 result in hyperactivation of p110δ and thus increase activation of the PI3K signalling pathway and its downstream effectors. Due to the involvement of PI3K in multiple immune cell functions, APDS1 is a condition marked by a varied array of clinical manifestations, including hepatosplenomegaly, lymphoproliferation, poor responses to vaccines, susceptibility to herpesvirus infections, cytopenias and B-cell lymphomas [1–3, 6]. Along with the lymphoproliferation and splenomegaly seen in APDS1 patients, there is also increased T cell activation, with decreased naïve cells and increased memory cells observed in both the CD4^+^ and CD8^+^ compartments [1, 2, 6–13].

Previous work in both APDS1 patients and mouse models of APDS1 (that express the *Pik3cd* E1020K GOF variant analogous to the E1021K variant found in the majority of patients), showed that T cell function is substantially altered by increased PI3K signalling. First, CD4^+^T cells are skewed towards a Th1 and Th2 phenotype. Second, the Tfh population is increased in frequency, however, these cells have decreased B-cell helper function, which contributes to the poor antibody response seen in APDS1 patients [6, 13–15]. Third, while the Treg frequency (CD4^+^CD127^lo^CD25^hi^ or CD4^+^FoxP3^+^CD25^+^) in *PIK3CD* GOF patients is equivalent to healthy controls [14, 16], the Treg population in *Pik3cd* GOF mice is significantly increased when defined by FoxP3 expression [14]. Lastly, PI3K^GOF^ CD8^+^ T cells have an exhausted phenotype, undergo increased cell death and exhibit metabolic fatigue, contributing to the recurrent EBV and/or cytomegalovirus infection [6–8, 17]. While our understanding of the effects of the PI3K GOF on T cell biology has expanded dramatically over the last decade it is still unclear which signals promote the T cell activation seen in APDS1 patients and cause the decrease in naïve T cells. Our previous work [14] showed that T cell activation in PI3K^GOF^ is extrinsically driven, as WT T cells that developed in the presence of PI3K GOF cells in mixed WT:PI3K GOF bone marrow chimeras exhibited increased levels of activation, including increased WT Tfh and Tregs [14].

Here, we sought to understand what the main extrinsic drivers are that promote and sustain CD4^+^ T cell activation and altered differentiation in APDS1 patients. To avoid complicating factors of chronic or recurrent infection that may impact T cell activation in patients we used our *Pik3cd* E1020K GOF mouse model of APDS1[11] (hereafter referred to as PI3K^GOF^). We used multiple mixed bone marrow chimera strategies to specifically evaluate the contribution of different immune cells to the increased T cell activation and augmented Tfh and Treg differentiation. Our findings reveal that T cells themselves are the main contributors to modulating the environment to promote and sustain increased CD4^+^ T cell activation. In contrast, expansion of Tfh cells is driven by PI3K GOF B cells. Lastly, we demonstrate that the dysregulated Treg compartment is impacted by both intrinsic factors and extrinsic influences from multiple cell types. Importantly, however PI3K GOF Tregs did not have an increased propensity to form ex-Tregs or inflammatory Tregs.

## Results

### The PI3K GOF environment drives hyperactivation and differentiation of WT CD4^+^ T cells

We and others have previously reported that T cell activation due to PI3K GOF is driven primarily by T cell extrinsic mechanisms [14, 17, 18]. We confirmed those findings here by generating 50:50 mixed BM chimeras (WT:WT or WT:PI3K^GOF^ – Fig 1A). As shown previously, there was a pronounced decrease in the frequency of naïve cells (Fig 1B) and corresponding increase in central memory (Fig 1C) and effector memory (Fig 1D) cells within the WT CD4^+^T cell population that developed in the presence of PI3K GOF cells (red bar on the left) compared to those developing in the presence of WT cells (black bar on the left). We also observed a significant increase in the frequencies of WT Tfh (Fig1E) and WT Tregs (Fig 1F) in mixed WT:PI3K GOF chimeras. Similar trends in the activation profile of CD8^+^T cells were also observed (data not shown). Altogether, the activated phenotype observed in the WT T cells in WT:PI3K GOF mixed chimeras indicates that the PI3K GOF environment is able to extrinsically drive T cell activation.

**Fig 1.**
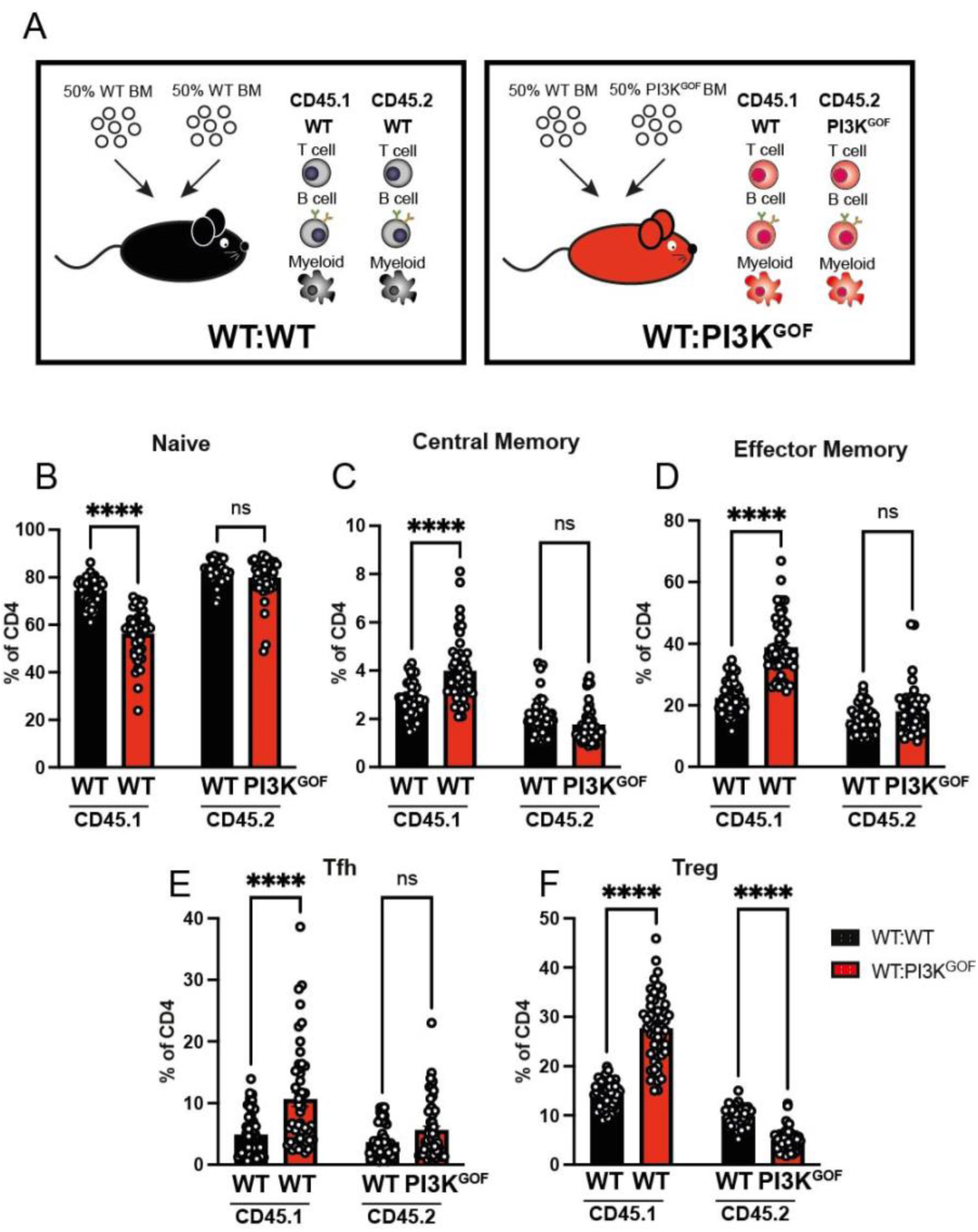
PI3K GOF cells drive hyperactivation and differentiation of WT CD4^+^ T cells through extrinsic mechanisms. **A**, WT:WT or WT:PI3KGOF mixed BM chimeras were generated as shown in the schematics. **B-F**, Spleens from 11-15 weeks after reconstitution were stained to identify different CD4^+^ T cell populations within the CD45.1^+^ or CD45.2^+^ compartments. Percentages of the CD4^+^ T cell compartment in each of the chimeric combination gating for naive (CD44^lo^CD62L^hi^), central memory (CD44^hi^CD62L^hi^), effector memory (CD44^hi^CD62L^lo^), Tfh (CXCR5^+^PD1^+^) and Treg (FoxP3^+^) CD4^+^ T cells were determined (Each point represents CD45.1^+^ or CD45.2^+^ cells in a different mouse, bars show means ± SEM, n=49-51). Significant differences were determined by 2way ANOVA: ****P <0.0001, ns – not significant.

### CD4^+^T cell activation is driven by PI3K GOF T and/or B cells

To investigate the extrinsic driver of PI3K-mediated CD4^+^ T cell activation and altered differentiation, we generated mixed bone marrow chimeras in which we excluded certain cell types to determine their specific contribution towards the T cell activation phenotype. We started by crossing PI3K^GOF^ mice with RAG^KO^ mice to generate PI3K^GOF^RAG^KO^ mice which cannot generate T or B cells [19]. We then generated 50:50 mixed bone marrow chimeras (WT:WT, WT:PI3K^GOF^, WT:WT RAG^KO^, and WT:PI3K^GOF^ RAG^KO^ – Fig 2A) to delineate the contribution of PI3K GOF T and B cells versus other non-T/non-B hematopoietic cells (e.g. myeloid cells, NK cells) to the activation and differentiation of WT CD4^+^ T cells. Thus, the bars shown in Figure 2 represent exclusively the WT CD4 T cells and how the PI3K hematopoietic cells impacted them. In the absence of PI3K GOF B and T lymphocytes, the WT CD4^+^ T cell population (Fig 2B, light green bar) did not show the same decrease in naïve cells and increase in memory cells that developed in the presence of PI3K GOF cells of all lineages (Fig 2B-D, red bar). These data suggest that PI3K GOF B and/or T cells are key drivers of the activated CD4^+^T cell phenotype we observe in APDS1 patients and mice. However, removal of T and B cells in WT:PI3K^GOF^RAG^KO^ chimeras did not completely ablate changes in WT CD4^+^ T cells. In fact, we observed modest but significant decreases in naïve cell and increases in effector memory cells compared to the WT:WT RAG^KO^ control chimeras (Fig 2D, dark green), suggesting that other cells such as myeloid cells may play a role.

**Fig 2.**
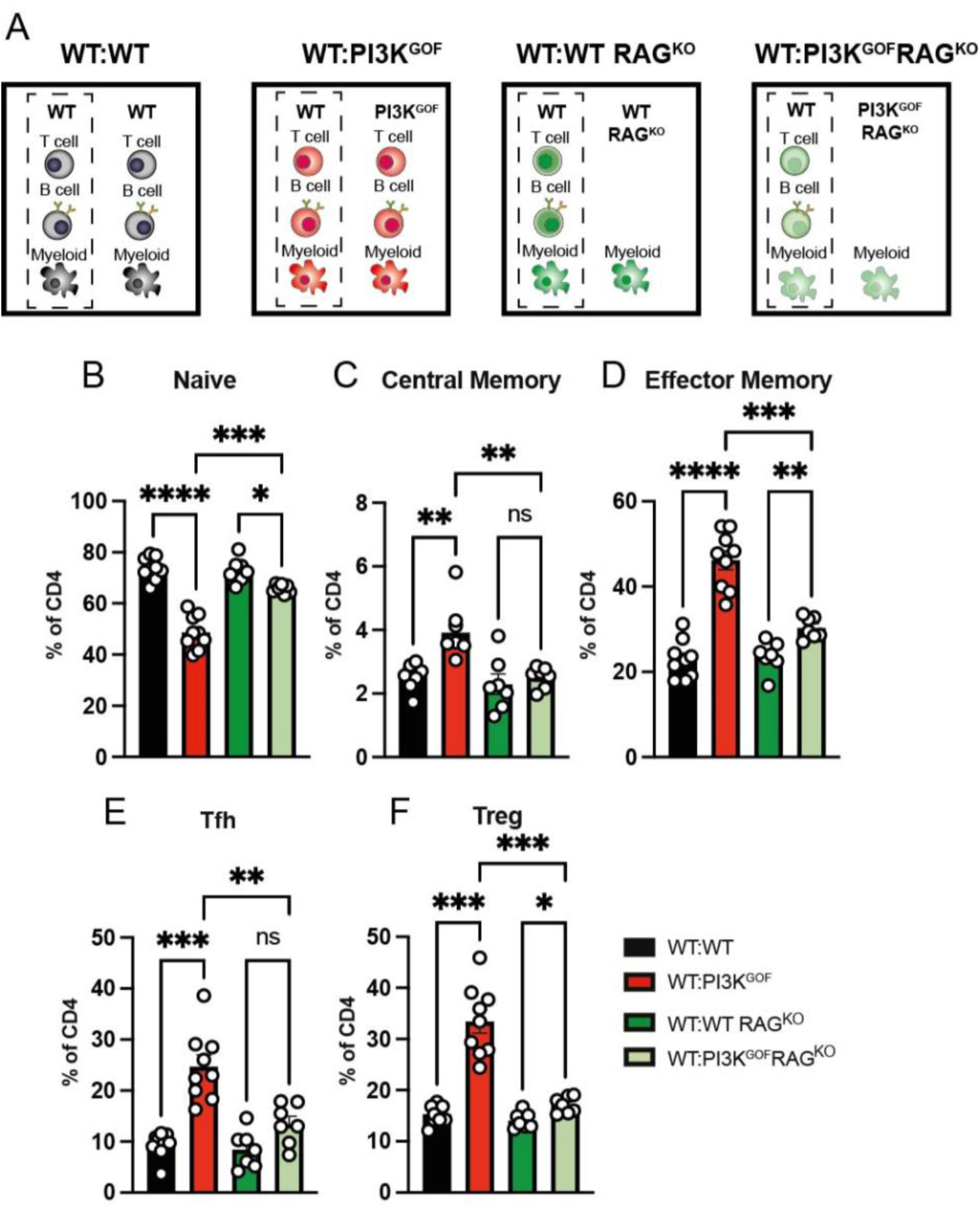
PI3K GOF non-T/non-B cells exert mild effect driving T cell hyperactivation or Tfh and Treg differentiation. **A**, WT:WT, WT:PI3KGOF, WT:WT RAGKO and WT:PI3K^GOF^RAGKO mixed BM chimeras were generated as shown in schematics. **B-F**, Spleens from 13-15 weeks after reconstitution were stained to identify different CD4^+^ T cell populations within the CD45.1^+^ or CD45.2^+^ compartments. The percentage of naive (CD44^lo^CD62L^hi^), central memory (CD44^hi^CD62L^hi^), effector memory (CD44^hi^CD62L^lo^), Tfh (CXCR5^+^PD1^+^) and Treg (FoxP3^+^) CD4^+^ T cells (each point represents a different mouse, bars show mean ± SEM, n=13) in each chimeric combination was determined. For simplicity, only the WT cells in each chimera (indicated in the dashed box in Fig 2A) are shown. Significant differences were determined by using Brown-Forsythe and Welch ANOVA tests: *P <0.05, **P <0.01, ***P <0.001, and ****P <0.0001.

We also assessed if the PI3K GOF B and T cells altered the CD4^+^T cell differentiation towards Tfh and Tregs. We observed that in the absence of PI3K GOF T and B cells there was no increase in Tfh frequency (Fig 2E). The expansion of the WT Treg population was also greatly reduced in the absence of PI3K GOF T and B cells (Fig 2F), although there was still a significant albeit mild increase compared to the WT:WT RAG^KO^ control chimeras. This indicates that PI3K GOF T and/or B cells are critical for inducing the expansion of both Tfh cells and Treg cells.

### PI3K GOF T cells drive the activation of WT T cells

Having established that PI3K GOF B and/or T cells were the predominant cell type inducing changes observed in the CD4^+^ T cells, we sought to unravel the contribution of each of these cell types. To assess the role of T cells, we crossed PI3K^GOF^ mice with CD3^KO^mice, generating a PI3K^GOF^CD3^KO^ line, in which the mice have all the PI3K GOF cells except T cells. We then generated 50:50 mixed bone marrow chimeras (WT:WT, WT:PI3K^GOF^, WT:WT.CD3^KO^, and WT:PI3K^GOF^CD3^KO^ – Fig 3A), so we could assess the contribution of PI3K GOF T cells to driving WT CD4^+^T cell activation. We observed that in the WT:PI3K^GOF^CD3^KO^ chimeras, which lacked PI3K GOF T cells, the frequencies of naïve, central memory and effector memory WT CD4^+^T cells were almost restored to levels observed in control chimeras (Fig 3B-D). These results reveal the importance of the PI3K GOF T cells in modulating the environment and promoting exacerbated T cell activation, which is a signature clinical feature observed in APDS1 patients.

**Fig 3.**
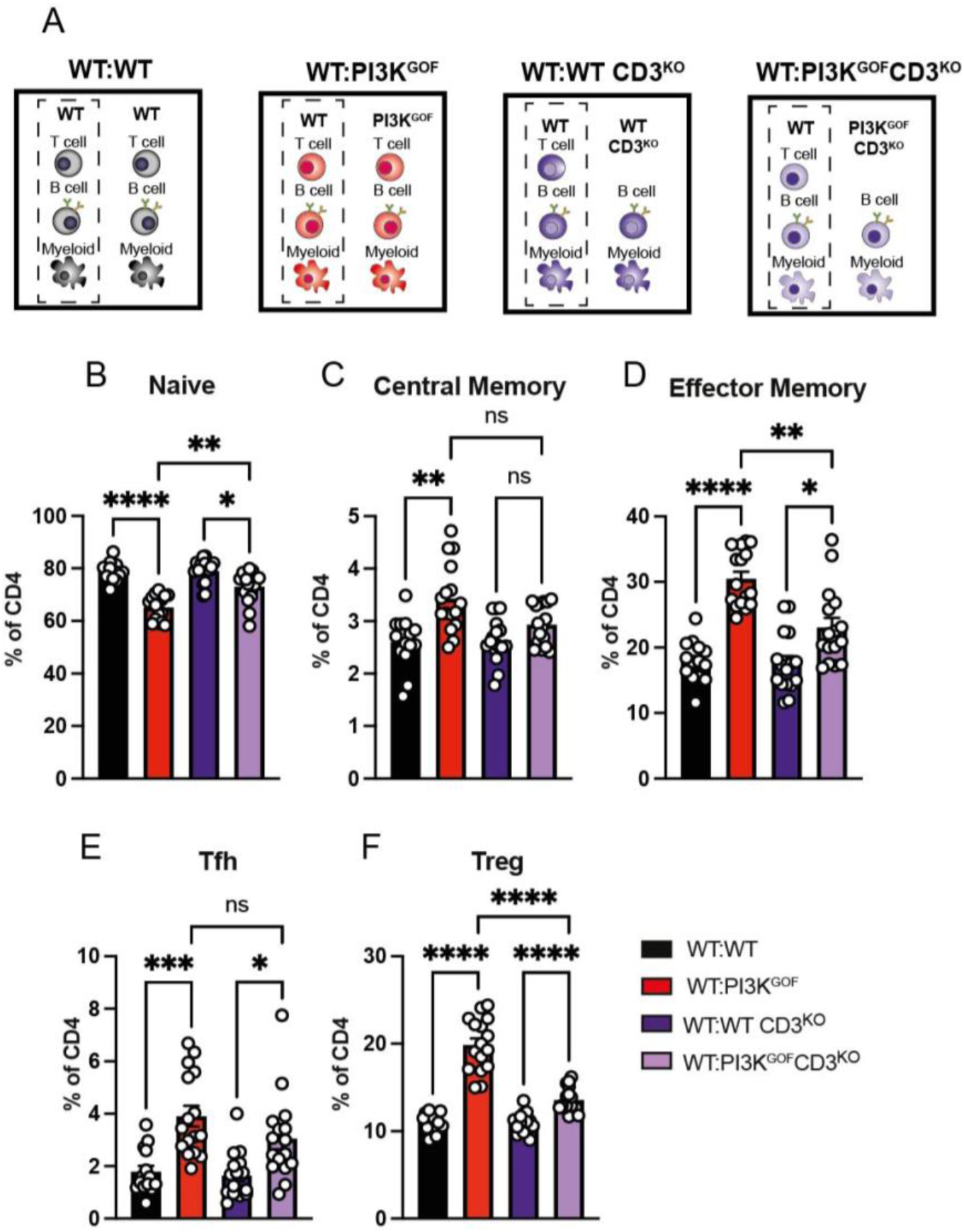
PI3K GOF T cells drive WT CD4^+^ T cell hyperactivation. **A**, WT:WT, WT:PI3K^GOF^, WT:WT CD3KO and WT:PI3K^GOF^CD3^KO^ mixed BM chimeras were generated as shown in the schematics. **B-F**, Spleens from 12-14 weeks after reconstitution were stained to identify different CD4^+^ T cell populations within the CD45.1^+^ or CD45.2^+^ compartments. Percentages of the WT CD4^+^ T cell compartment (shown in dashed boxes in A) in each of the chimeric combination gating for naive (CD44^lo^CD62L^hi^), central memory (CD44^hi^CD62L^hi^), effector memory (CD44^hi^CD62L^lo^), Tfh (CXCR5^+^PD1^+^) and Treg (FoxP3^+^) CD4^+^ T cells were determined (each point represents a different mouse, bars show mean ± SEM, n=14-16). Significant differences were determined by using Brown-Forsythe and Welch ANOVA tests: *P <0.05, **P <.01, ***P <0.001, and ****P <0.0001.

Given the critical role of PI3K GOF T cells in driving T cell activation and increases in memory cells we were also interested in assessing whether PI3K GOF T cells contributed to the dysregulation of Tfh and Treg cells. In contrast to what was seen with memory T cells, we observed no significant decrease in the Tfh frequency in the absence of PI3K GOF T cells in the WT:PI3K^GOF^CD3^KO^ chimeras (Fig 3E), arguing against the PI3K GOF T cells being the main cellular mediator of Tfh expansion. Next, we quantified Treg frequencies and observed that the expansion of WT Tregs that is observed in the WT:PI3K^GOF^ chimeras is significantly blunted in the absence of PI3K GOF T cells in the WT:PI3K^GOF^CD3^KO^ chimeras (Fig 3F), highlighting an important role of the PI3K GOF T cells in modulating the immune environment and triggering Treg expansion.

### PI3K GOF Tregs support the expansion of WT Tregs but not the overall CD4^+^ T cell activation

Having established that T cells were the PI3K GOF cell responsible for inducing most of the WT T cell activation observed, we sought to dissect which PI3K GOF T cell population was responsible for this effect. Several lines of evidence suggested that dysregulated Tregs driven by overactive PI3K might play a role. First, we previously showed that PI3K GOF causes an increase in CD25^lo^ Tregs [14]. Second, other models of PI3K overactivation, such as PTEN deficiency, show dysregulation of Tregs. PTEN counteracts the actions of PI3K by promoting the dephosphorylation of PIP_3_ back into PIP_2_ [4], thus PTEN deficiency leads to an increase in PIP_3_ and downstream signalling. It has previously been shown that in mice in which PTEN is deleted in Tregs (PTEN^ΔTreg^), there is an accumulation of an “ex-Treg” population [20, 21]. Ex-Tregs are cells that previously expressed FoxP3, the master regulator of the Tregs, but went through a remodulation process that results in FoxP3 downregulation. This was associated with not only a loss of the regulatory ability but also the acquisition of a pro-inflammatory signature, such as expression of IFNγ [20, 21]. As a result, these PTEN^ΔTreg^ mice that accumulated ex-Tregs have increased overall T cell activation. Considering the parallel between the PTEN^ΔTreg^ mice and our PI3K^GOF^ mouse model, and the pronounced increase in CD25^−^ Treg we have observed in aged PI3K GOF Treg mice [14], we hypothesised that pathogenic ex-Tregs may also explain the increased CD4^+^ T cell activation. To investigate this, we crossed FoxP3^GFP-DTR^ mice [22], in which GFP (green fluorescent protein) and DTR (diphtheria toxin receptor) are expressed from the *Foxp3* locus, to PI3K^GOF^ mice to generate a PI3K^GOF^FoxP3^GFP-DTR^ line. This allowed us to sort Tregs based on GFP expression as well as delete them through a course of diphtheria toxin (DT) injections. We generated 50:50 mixed bone marrow chimeras, but this time using PI3K^GOF^FoxP3^GFP-DTR^ mice (WT:WT, WT:PI3K^GOF^, WT:WT FoxP3^GFP-DTR^, and WT:PI3K^GOF^FoxP3^GFP-DTR^ – Fig 4A). Three weeks post irradiation and reconstitution, these chimeras underwent a Treg depletion regimen (Fig 4B) involving intravenous injections of DT every two weeks until their analysis at week 11. This resulted in sustained depletion of the FoxP3^GFP-DTR^ cells, while FoxP3^WT^ Tregs that did not express the DTR transgene were not affected by the treatment, as murine cells do not endogenously express this receptor (Supplementary Fig 1). This system ensured that the CD45.1^+^ WT compartment of both WT:WT FoxP3^GFP-DTR^ and WT:PI3K^GOF^FoxP3^GFP-DTR^ chimeras maintained functional Tregs, ostensibly preventing development of the severe autoimmune phenotype observed in mice that lack Tregs [23]. This allowed us to assess the specific contribution of the PI3K GOF Tregs in driving the T cell activation and altered differentiation.

**Fig 4.**
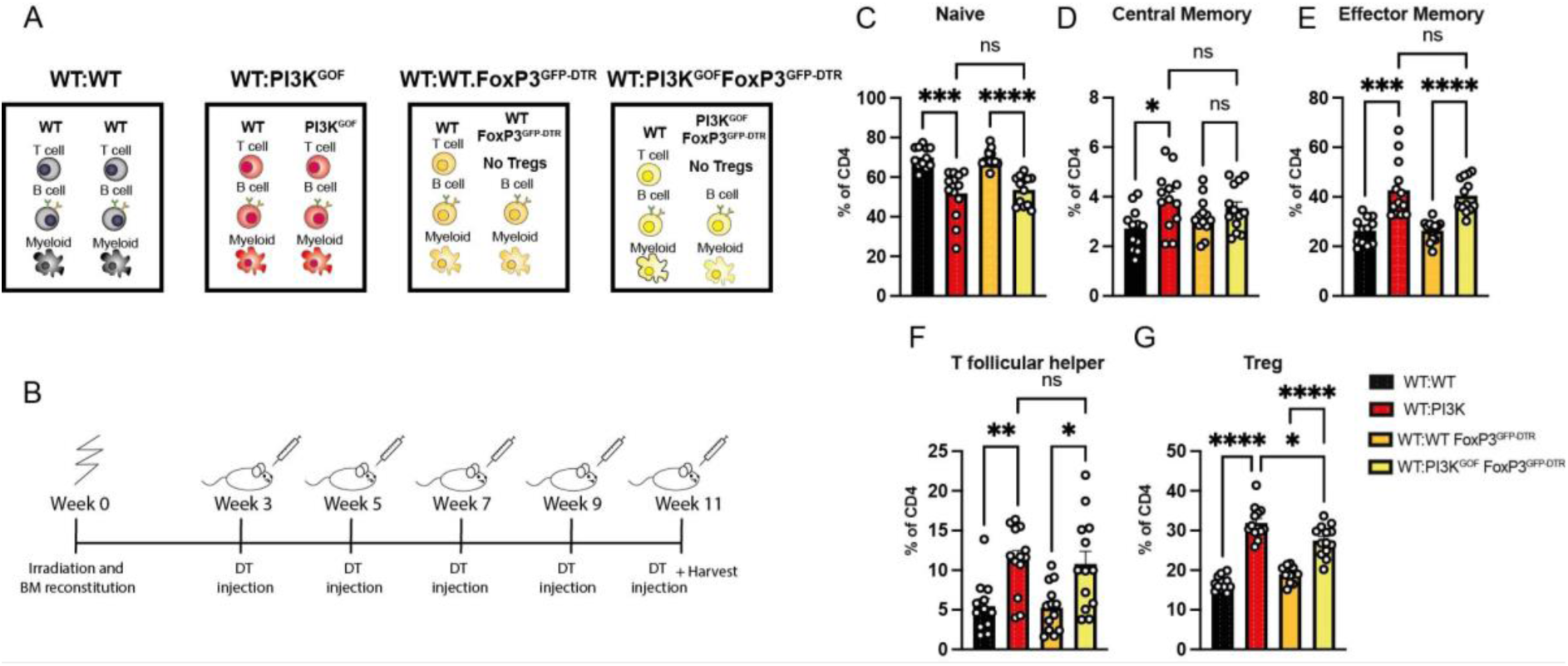
PI3K GOF Tregs cells are not the drivers of CD4^+^ T cell hyperactivation. **A**, WT:WT, WT:PI3K^GOF^, WT:WT FoxP3^GFP-DTR^ and WT:PI3K^GOF^FoxP3^GFP-DTR^ mixed BM chimeras were generated as shown in the schematics. **B**, Schematics of diphtheria toxin injection regimen to achieve Treg depletion. **C-G**, Spleens from stained to identify different CD4^+^ T cell populations within the CD45.1^+^ or CD45.2^+^ compartments 11 weeks after reconstitution and diphtheria toxin Treg depletion regimen. Percentages of the WT CD4^+^ T cell compartment in each of the chimeric combination gating for naive (CD44^lo^CD62L^hi^), central memory (CD44^hi^CD62L^hi^), effector memory (CD44^hi^CD62L^lo^), Tfh (CXCR5^+^PD1^+^) and Treg (FoxP3^+^) CD4^+^ T cells were determined (each point represents a different mouse, bars show means ± SEM, n=12-13). Significant differences were determined by using Brown-Forsythe and Welch ANOVA tests: *P <0.05,**P <0.01,***P <0.001, and ****P <0.0001.

The proportions of naïve and memory CD4+ T cells detected in the WT:PI3K GOF chimeras were similar irrespective of the presence or absence of PI3K GOF Tregs (Fig 4C-E; light yellow vs red bars). Similarly, we saw no effect of depletion of PI3K GOF Tregs on the increase in Tfh cells (Fig 4F). However, there was a small but significant decrease in the expansion of WT Treg in the absence of PI3K GOF Tregs (Fig 4G). Thus, in contrast to previous observations in the PTEN^ΔTreg^ mice [20, 21], our results suggest that the PI3K GOF Tregs are not responsible for promoting a pro-inflammatory environment that drives enhanced CD4^+^ T cell activation and increased memory and Tfh cells.

This raised the question of whether PI3K GOF T regs have the same propensity as PTEN^ΔTreg^ to become ex-Tregs. To explore this, we sorted either WT or PI3K GOF GFP-expressing cells (CD4^+^FoxP3^GFP+^) and transferred them into congenic WT mice (Fig 5A). When we analysed these mice 4 weeks later, we observed a reduced frequency of donor Tregs in mice that received PI3K GOF cells when compared to mice that received WT Tregs (Fig 5B), suggesting PI3K GOF Tregs have reduced survival. Within the donor population, we also defined Tregs (CD4^+^GFP^+^FoxP3^+^) and ex-Tregs (CD4^+^GFP^−^FoxP3^−^). Surprisingly, despite previous reports of the critical role of regulated PI3K signalling in maintaining Treg identity, we did not observe any differences in the frequency of ex-Tregs (Fig 5C) and normal Tregs (Fig 5D) between WT and PI3K GOF transferred Tregs. These results suggest that, at least in our system, PI3K GOF due to a *Pik3cd* GOF variant does not drive an increase in the development of ex-Tregs.

**Figure 5.**
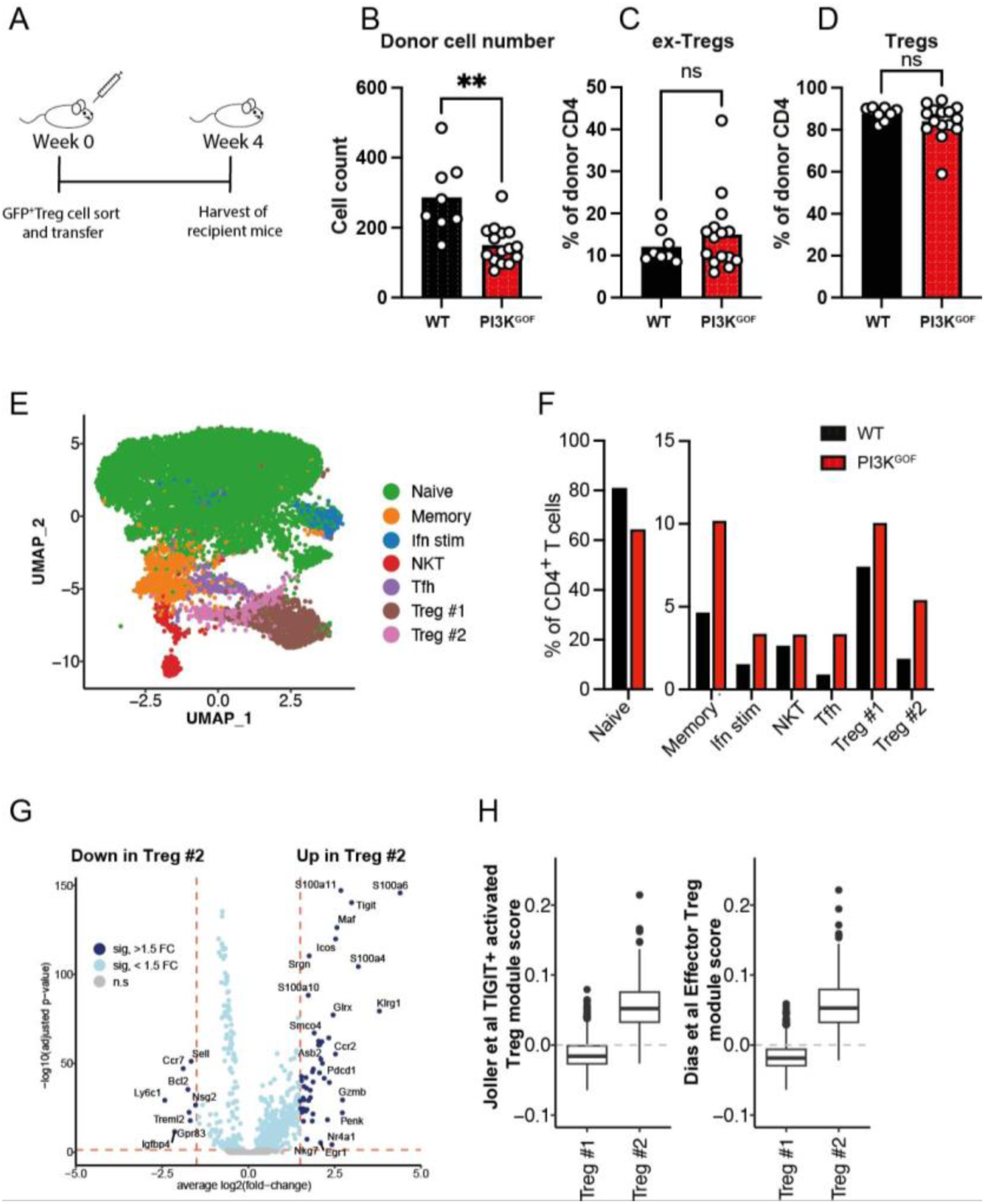
Pik3cd GOF does not induce inflammatory ex-Treg phenotype. **A**, Schematics of Treg adoptive transfer **B**, Donor cell number following 4 weeks of adoptive transfer **C-D**, Percentages of ex-Tregs (donor CD4^+^FoxP3^−^GFP^−^) and Tregs (donor CD4^+^FoxP3^+^GFP^+^) (each point represents a different mouse, bars show means, n=8-15). Significant differences were determined by unpaired Students t-test, *P<0.05, **P<0.01; ****P<0.0001. **E-H,** CD4^+^ T cells were isolated from WT (n=2) and PI3K GOF (n=2) spleens and scRNA-seq performed. **E**, UMAP shows clustering based on all cells; **F**, Percentage of cells in each cluster. **G**, differentially expressed (DE) genes between Treg #2 and Treg #1. **H**, Enrichment of DE genes against effector Treg data sets from Joller et al [25] and Dias et al [24].

To explore this further, we performed scRNA-seq on CD4^+^ T cells isolated from WT or PI3K GOF mice and identified 7 different populations (Fig 5E). Consistent with our flow cytometry data, scRNA-seq data revealed a loss of naïve CD4^+^ T cells in PI3K GOF mice together with an increase in Tfh and Treg cells (Fig 5F). Interestingly clustering of the data revealed two distinct Treg populations, with one population (Treg #2) particularly enriched in PI3K GOF mice (Fig 5E,F). This population showed increased expression of genes such as *Icos*, *Pdcd1, Tigit, Cd44* and *Ctla4* and decreased expression of *Ccr7* and *Sell* (encoding CD62L) (Fig 5G) consistent with an effector Treg phenotype [24, 25]. Interestingly the Treg #2 population also expressed high levels of *Klrg1* which has been associated with short-lived effector Tregs [26], providing a potential explanation for the decreased survival of transferred Tregs. In contrast, they did not show increased expression of genes such as *Bcl6* and *Il21* which were associated with ex-Tregs in the absence of Pten [21]. Taken together, these experiments demonstrate that *Pik3cd* GOF does not lead to destabilisation of the Treg phenotype and an increase in ex-Tregs but rather leads to an increase in effector Tregs.

### PI3K GOF B cells drive the expansion of Tfh cells but not of memory CD4^+^ T cells

Our data showed that in the absence of PI3K GOF T cells many of the changes induced in the WT CD4^+^ T cells that developed in mixed chimeras were rescued (Fig 3). However, some changes, such as the increased Tfh population, were not rescued by the absence of T cells. Thus, we tested whether PI3K GOF B cells played a role in the phenotype induced in CD4^+^ T cells, particularly the induction of Tfh cells. For this, we crossed the PI3K^GOF^ mice with Cd79a^KO^ mice, which lack B cells, and generated 50:50 mixed bone marrow chimeras (WT:WT, WT:PI3K^GOF^, WT:WT Cd79a^KO^, and WT:PI3K^GOF^Cd79a^KO^ – Fig 6A). The absence of PI3K GOF B cells did not rescue the decrease in naïve WT T cells nor the increase in memory WT T cells, suggesting that PI3K GOF B cells do not drive this phenotype (Fig 6B-D). However, in the absence of PI3K GOF B cells, the WT Tfh frequencies were restored to normal levels (Fig 6E – light blue column). These results suggest that PI3K GOF induces intrinsic dysregulation of B cells which in turn stimulates an expansion of Tfh cells. Consistent with this, we found that that PI3K GOF B cells mixed bone marrow chimeras have increased expression of the costimulatory molecules CD80 and CD86 and a higher proportion of cells with a GC phenotype (Supp Fig 2), indicating a cell intrinsic dysregulation of B cells.

**Fig 6.**
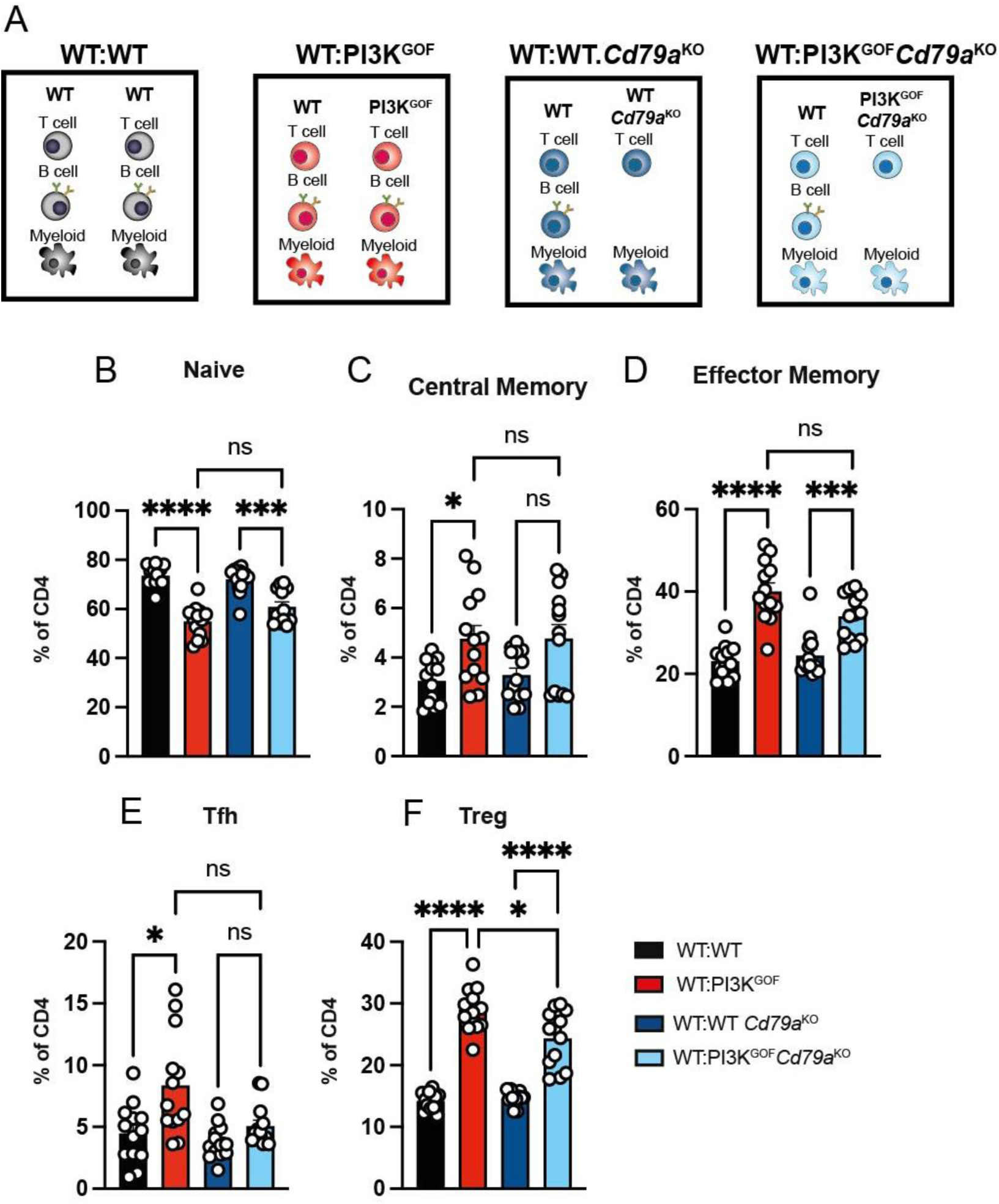
PI3K GOF B cells drive increased Tfh differentiation. **A**, WT:WT, WT:PI3K^GOF^, WT:WT Cd79a^KO^ and WT:PI3K^GOF^Cd79a^KO^ mixed BM chimeras were generated as shown in the schematics. **B-F**, Spleens from 13-14 weeks after reconstitution were stained to identify different CD4^+^ T cell populations within the CD45.1^+^ or CD45.2^+^ compartments. Percentages of the WT CD4+ T cell compartment in each of the chimeric combination gating for naive (CD44^lo^CD62L^hi^), central memory (CD44^hi^CD62L^hi^), effector memory (CD44^hi^CD62L^lo^), Tfh (CXCR5^+^PD1^+^) and Treg (FoxP3^+^) CD4^+^ T cells were determined (each point represents a different mouse, bars show mean ± SEM, n=13). Significant differences were determined by using Brown-Forsythe and Welch ANOVA tests: *P <0.05, ***P <0.001, and ****P <0.0001.

## Discussion

Patients with APDS1 show dramatic dysregulation of the T cell compartment with increased memory and Tfh cells, and lymphoproliferation [1, 2, 6–13]. Here we sought to elucidate cellular mechanisms underpinning the dysregulated T cell activation and differentiation that are observed in these patients. Our mouse model enabled us to determine the contribution of different immune cell populations and exclude environmental factors in patients such as recurrent or chronic infection, or any treatment regimes. This revealed that the T cell activation, characterised by a loss of naïve T cells and increase in memory T cells, can largely be driven in a cell extrinsic manner by PI3K GOF T cells that change the environment in such a way that even WT T cells take on an activated phenotype. Earlier studies that have suggested a critical role of PI3K in Treg function led us to examine whether this may be induced by dysregulated Tregs, that take on an inflammatory phenotype.

It has previously been demonstrated that deletion of *Pten* in Tregs using FoxP3-cre (PTEN^ΔTreg^) resulted in Treg instability, which in turn caused an expansion of inflammatory Tregs, that had lost CD25 expression and drove an increase in GC and Tfh cells [20, 21]. Given the similar phenotype of PTEN^ΔTreg^ and *Pik3cd* GOF mice and the increased PI3K signalling in both models, we hypothesised that *Pik3cd* GOF may have a similar effect. Surprisingly, our results revealed that in contrast to FoxP3-cre mediated loss of PTEN in Tregs, Tregs that developed in *Pik3cd* GOF mice did not show a propensity to become inflammatory ex-Tregs but instead maintained FoxP3 expression and displayed an effector phenotype. Further, we did not observe a role for *Pik3cd* GOF Tregs in promoting either T cell activation or Tfh generation. The reason for this difference between the PTEN^ΔTreg^ and the *Pik3cd* GOF Tregs is not clear but may reflect the timing and/or magnitude of increased PI3K signalling in the two models [27]. Indeed, while this manuscript was in preparation Singh et al reported findings from a mouse model whereby the *Pik3cd* E1020K GOF variant was specifically expressed in Tregs using the same FoxP3-cre (FoxP3-*Pik3cd* GOF)[28]. Similar to our results here, these investigators saw no evidence for *Pik3cd* GOF inducing ex-Tregs, suggesting that the nature or magnitude of the altered signalling induced by PI3Kd GOF is not as severe as that induced by complete PTEN deletion and that the T cell dysregulation in APDS1 patients is not caused by inflammatory ex-Tregs.

In contrast to our finding, however, Singh et al did observe increased GC formation and T cell activation in aged FoxP3-*Pik3cd* GOF mice. The authors suggested that this may be driven by loss of function in FoxP3-*Pik3cd* GOF Tregs. However, this is not supported by the demonstration that FoxP3-*Pik3cd* GOF Tregs retain normal suppressive ability *in vitro* [28] and our scRNA-seq data showing Tregs in *Pik3cd* GOF mice have a transcriptome associated with increased effector function. Further, our results demonstrate that an increase in T cell activation persisted when *Pik3cd* GOF Tregs were depleted (Fig 4) demonstrating a clear role for non-Tregs in driving this process. It is possible that the different effects seen in FoxP3-*Pik3cd* GOF mice were driven by expression of the *Pik3cd* GOF allele in some B and non-Treg T cells as the FoxP3-cre employed by Singh et al has previously been shown to be leaky, resulting in recombination in small numbers of both non-Treg T cells and B cells [27, 29]. However, we cannot rule out that *Pik3cd* GOF Tregs have mildly reduced suppressive ability such that over time a lack of regulation could contribute to T cell activation in the host which was not detected in our shorter time frames (11-15 weeks).

In contrast to the key role of *Pik3cd* GOF T cells in driving the general T cell activation and loss of naïve T cells, our results clearly demonstrated that PI3K GOF B cells were responsible for driving increased Tfh cells in the mice. We also observed that PI3K GOF B cells expressed higher levels of costimulatory molecules such as CD80/CD86 and formed more GCs. This is consistent with increased and/or dysregulated GCs observed in APDS1 patients and PI3K GOF mice [2, 3, 13, 30–32] and our previous findings that increased PI3K due to a *Pik3cd* GOF variant drives a B cell intrinsic loss of tolerance and the generation of spontaneous GCs [30]. Together these results suggest that the increased Tfh cells may be driven by autoreactive B cells that inappropriately present antigen and activate CD4^+^ T cells. It would be interesting in the future to assess the antigen specificity of Tfh and GC B cells in *Pik3cd* GOF mice and APDS patients to confirm whether they are indeed autoreactive.

## Methods

### Mice

All experiments were approved by the Garvan Institute–St. Vincent’s Animal Ethics Committee. All mice were bred and housed in specific pathogen free (SPF) conditions at Australian BioResources (ABR – Moss Vale) and the Garvan Institute Biological Testing Facility. *Pik3cd*^E1020K^ mice (referred to as PI3K^GOF^ mice) have been previously described [11]. These mice are heterozygous for a G to A base substitution, resulting in a Glu-to-Lys amino acid substitution in p110δ at amino acid residue 1020 (i.e. E1020K; corresponding to E1021K in human p110δ, the most common variant identified in APDS1 patients). Multiple crosses were made at ABR as follows: PI3K^GOF^ mice were crossed with *Rag^−/-^* (CD45.1 congenic – Ptprc^a/a^ background), resulting in the PI3K^GOF^RAG^KO^CD45.1^+^ line; PI3K^GOF^ mice were crossed with CD3e^KO^ mice [33], resulting in the PI3K^GOF^CD3^KO^ line; PI3K^GOF^ mice were crossed with *Cd79a*-deficient mice [34], resulting in the PI3K^GOF^*Cd79a*^KO^ line; PI3K^GOF^ mice were crossed with *FoxP3*^GFP-DTR^ mice [22], resulting in the PI3K^GOF^*FoxP3*^GFP-DTR^ line. All mice were 6-12 weeks old at the start of experiments and were age and sex-matched.

### Mixed bone marrow chimeras

For mixed bone marrow (BM) chimera experiments, donor cells were harvested and processed. Recipient mice were sub-lethally irradiated twice at 475 rad 6 hours apart. A 50:50 mix of each donor combination was injected intravenously (2-5×10^6^ cells/mouse) into recipient mice 6-12 weeks old C57Bl/6 (CD45.1 or CD45.2 congenic) purchased from Australian BioResources. For the *FoxP3*^GFP-DTR^ chimeras, controls and *FoxP3*^GFP-DTR^ groups received 50ng/mg intravenously of diphtheria toxin (DT) every fortnight. All chimeras were harvested 11-15 weeks post reconstitution.

### Adoptive transfer

For adoptive transfer experiments, donor spleens (WT *FoxP3*^GFP-DTR^ or PI3K^GOF^ *FoxP3*^GFP-DTR^) were harvested and processed. Splenocytes were enriched with the MACS CD4^+^ T-cell isolation kit (Miltenyi Biotech) and separated with either the AutoMACS Pro separator or manual columns. Recovered cells were cell sorted on a FACSAria (BD Biosciences) as CD4^+^GFP^+^ cells. 5×10^5^ sorted cells were adoptively transferred intravenously into 6-12 weeks old C57Bl/6 (CD45.1 congenic) purchased from Australian BioResources. Recipient mice were harvested 4 weeks following adoptive transfer and splenocytes were analysed.

### Flow cytometry reagents

The following reagents were purchased from BD Biosciences: anti-CD8a (53-6.7) Pacific Blue and BUV395, anti-CD16/CD32 (2.4G2) purified, anti-CD19 (1D3) BV510, anti-CD23(B3B4) BV605, anti-CD25(PC61) PE, anti-CD44 (IM7) BV605, anti-CD45.1(A20) BV421, anti-CD45.2 (104) BUV395, anti-CD45R/B220 (RA3-6B2) BV786, anti-CD95 (Jo2) PE, anti-CD138 (281-2) biotin, anti-CD185(CXCR5; 2G8) purified, anti-CD62L(MEL-14) APC, anti-IgM (AF6-78) PE, anti-PD1 (J43) PE, Streptavidin –BV605, –BUV395, –BV711. The following reagents were purchased form Biolegend: anti-CD21/34 (7E9) Pacific Blue, anti-CD24 (M1/69) Pacific Blue, anti-CD38 (90) PE-Cy7, anti-CD38 (90) APC-Cy7, anti-CD86 (GL-1) BV650, anti-CD93 (AA4.1) PerCP-Cy5.5, anti-IgD (11-26c.2a) APC-Cy7, anti-γδ TCR (GL3) PE-Cy7, Streptavidin BV421. The following reagents were purchased from eBioscience: anti-CD4 (RM4-5) APC-eFluor 780, anti-CD23 (B3B4) PE-Cy7, anti-CD25 (PC61.5) APC, anti-CD45.1 (A20) FITC, anti-CD45.1 (A20) PE-Cy7, anti-CD45.2 (104) APC-eFluor 780 and PerCP-Cy5.5, anti-CD62L (MEL-14) FITC, anti-FoxP3 (JK-15s) PE-Cy7, Streptavidin PE-Cy7

Normal mouse serum and Donkey anti-Rat IgG (H+L) Biotinylated were purchased from Jackson Immunoresearch. Normal Rat Serum was purchased from Sigma. The Foxp3/transcription factor staining buffer set (eBioscience/Life Technologies – Thermo Fisher) was used for the intracellular staining of FoxP3.

### scRNA-seq

Flow sorted CD4^+^ T cells from WT (n= 2) and PI3K^GOF^ (n= 2) mice were subject to scRNA-seq using the Chromium 5’ gene expression profiling (v2) from 10x Genomics. Totalseq-C hashtag antibodies (BioLegend) were used to multiplex the mice into a single 10x capture: C301ACCCACCAGTAAGAC (WT1), C302GGTCGAGAGCATTCA (WT2), C303CTTGCCGCATGTCAT (PI3K^GOF^1) and C304AAGCATTCTTCACG (PI3K^GOF^2). Gene expression libraries were processed using Cell Ranger (version 6.0.2) (10x Genomics) with default parameters and mm10-3.0.0 as reference. Hashtag libraries were processed with CITE-seq-count (version 1.4.5) [35] with the following parameters: –-start-trim 10 –-cbf 1 –-cbl 16 –-umif 17 –-umil 28. Hashtags were demultiplexed using Seurat’s (version 4.2.0) [36] HTODemux function.

Transcriptome libraries were first filtered to remove cells with high mitochondrial genes (>20%), low gene count and low library sizes using scater [37]. Low gene count and library sizes were determined using scater’s isOutlier with outliers defined as 2 median-absolute-deviations away from the median. Only barcodes with both gene expression and hashtag data were retained. Prior to creation of Seurat objects TCR V(D)J genes were removed from the count matrix to ensure that clustering wasn’t driven by T cell receptor gene usage. Gene count data was normalised using SCTransform. A Seurat workflow of RunPCA, RunUMAP, FindNeighbors and FindClusters was then applied. Gene expression between cell populations of interest was determined using the FindMarkers and FindAllMarkers functions.

Gene sets for Treg populations were scored using Seurat’s AddModuleScore with gene lists collected from Joller *et al.* [25] and Dias *et al.* [24].

### Statistical analysis

Significant differences were determined using Prism (GraphPad Software). Asterisks indicate statistical significance (*, P < 0.05; **, P < 0.01; ***, P < 0.001; ****, P < 0.0001).

## Acknowledgements

This work was supported by Leadership 2 Investigator Grant (2026131 to E.K. Deenick), project grant (1127157 to S.G. Tangye, E.K. Deenick), and a Leadership 3 Investigator Grant (1176665 to S.G. Tangye) awarded by the National Health and Medical Research Council. J. Bier was supported through the American Association of Immunologists Careers in Immunology Fellowship Program.

## Author contributions

J Bier, A Lau, KJL Jackson and S Ruiz-Diaz performed experiments and analysed data; R Brink generated mice; EK Deenick and SG Tangye provided supervision; EK Deenick designed and led the project; J Bier and EK Deenick interpreted the data and wrote the paper. All authors edited the manuscript.

