## Supplementary figures for "Both T and B cells contribute to dysregulated activation and differentiation of CD4^+^ T cells in Activated PI3K delta syndrome 1"

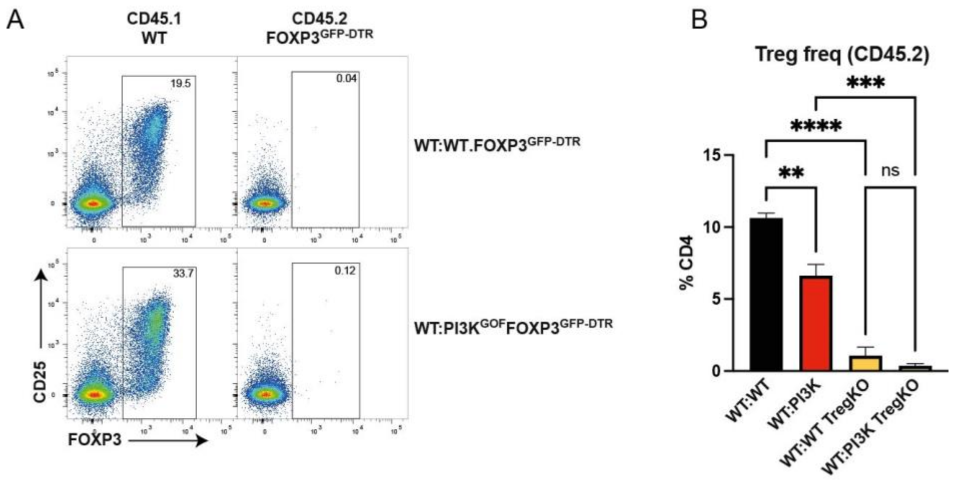


### Supplementary Figure 1. Depletion of FoxP3$^{GFP\text{-}DTR}$ Tregs

Mixed chimeras were set up as in Figure 4A and treated with diphtheria toxin as shown in Fig 4B. Spleens from mixed chimeras were stained to identify different CD4$^{+}$FoxP3$^{+}$ T cells within the CD45.1$^{+}$ or CD45.2$^{+}$ compartments. **A**, Representative flow plots are shown. **B**, Plot shows the number of FoxP3$^{+}$ cells within CD4 T cells in the CD45.2$^{+}$ compartment in each bone marrow chimera combination.

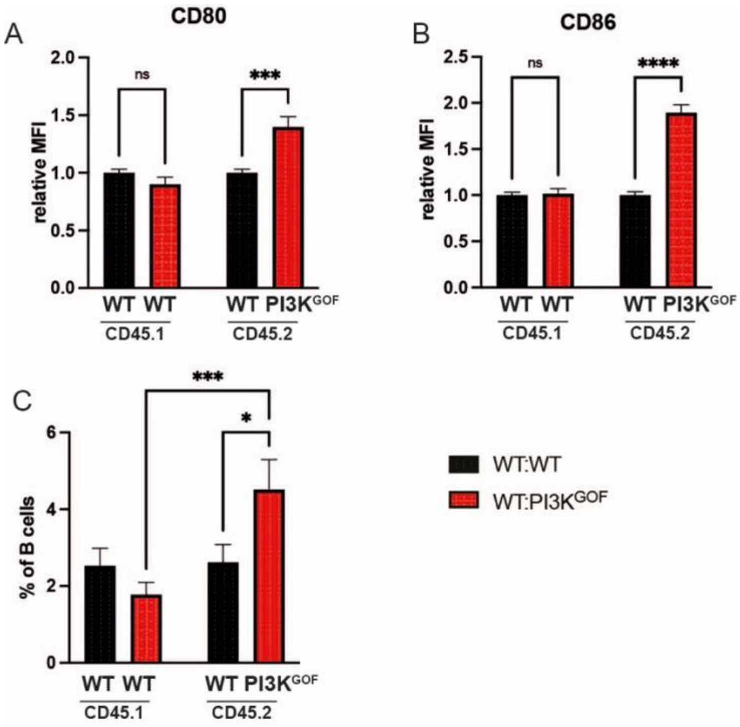


### Supplementary figure 2. PI3K GOF drives cell intrinsic activation of B cells

WT:WT or WT:PI3KGOF mixed BM chimeras were generated as shown in Fig 1A. Spleens from 11-15 weeks after reconstitution were stained to identify different B220$^+$ B cells within the CD45.1$^+$ or CD45.2$^+$ compartments. The expression of (**A**) CD80 and (**B**) CD86 was determined. Plots show MFI relative to WT cells in chimeras (mean ± SEM, n=5). **C**, the percentage of germinal centre (GC) B cells was determined (mean ± SEM, n=26-27).
